# Short and long-read genome sequencing methodologies for somatic variant detection; genomic analysis of a patient with diffuse large B-cell lymphoma

**DOI:** 10.1101/2020.03.24.999870

**Authors:** Hannah E Roberts, Maria Lopopolo, Alistair T Pagnamenta, Eshita Sharma, Duncan Parkes, Lorne Lonie, Colin Freeman, Samantha J L Knight, Gerton Lunter, Helene Dreau, Helen Lockstone, Jenny C Taylor, Anna Schuh, Rory Bowden, David Buck

**Author notes:** contributed equally (Hannah E Roberts, Maria Lopopolo, Alistair T Pagnamenta).

## Abstract

Recent advances in throughput and accuracy mean that the Oxford Nanopore Technologies (ONT) PromethION platform is a now a viable solution for genome sequencing. Much of the validation of bioinformatic tools for this long-read data has focussed on calling germline variants (including structural variants). Somatic variants are outnumbered many-fold by germline variants and their detection is further complicated by the effects of tumour purity/subclonality. Here, we evaluate the extent to which Nanopore sequencing enables genome-wide detection and analysis of somatic variation. We do this through sequencing tumour and germline genomes for a patient with diffuse B-cell lymphoma and comparing results with 150bp short-read sequencing of the same samples. Calling germline single nucleotide variants (SNVs) from the long-read data achieved good specificity and sensitivity. However, results of somatic SNV calling highlight the need for the development of specialized joint calling algorithms. We find the comparative performance of different tools varies significantly between structural variant types, and suggest long reads are especially advantageous for calling large somatic deletions and duplications. Finally, we highlight the utility of long reads for phasing clinically relevant variants, confirming that a somatic 1.6Mb deletion and a p.(Arg249Met) mutation involving *TP53* are oriented *in trans*.

## Introduction

Next generation sequencing (NGS) technologies have enabled a number of applications in genomics^1–5^. Whole Genome Sequencing (WGS) is among these and is the most comprehensive approach for characterising and analysing an individual’s genetic variation^6^. Substantial reductions in cost mean that WGS has become an increasingly important tool for clinical diagnosis and targeted treatment of rare disease and cancer^4,7–11^. Of particular importance to precision oncology is the ability of WGS to identify driver mutations (including those in non-coding regions)^12,13^, detect mutational signatures^14–16^, characterise structural variation and chromosomal rearrangements^17,18^, and pinpoint the genomic integration sites of oncoviruses, such as human papillomavirus^19^. However, the clinical interpretation of these results remains a challenge and precision oncology programmes and clinical trials involving NGS are underway^20^.

Although short-read, Illumina sequencing is considered the gold standard for the majority of clinical sequencing projects^21^, such data lead to biases even in WGS, due to uneven coverage of regions with high/low GC content and the difficulty of aligning short reads derived from repetitive DNA sequences^20,22,23^. Long read sequencing technologies, such as those developed by Pacific Biosciences (PacBio) and Oxford Nanopore Technologies (ONT), have proved invaluable for overcoming these challenges^24,25^. Thorough comparative studies have shown that long reads reduce the number of ‘ dark’ or ‘camouflaged’ regions of the genome^26^ and improve the sensitivity of structural variant (SV) detection^27^. Of course, both technologies have their pros and cons, but with fast turn-around times and lower start-up costs, and despite higher error rates of >10%, ONT WGS has already been used to resolve SVs in clinical cases^28,29^. Within cancer research specifically, low coverage ONT (Nanopore) WGS has been used for same-day diagnosis of brain tumours^30^, while targeted approaches have been developed for detecting *BCR-ABL1* fusion transcripts^31^, analysing prognostically relevant genes in chronic lymphocytic leukaemia^32,33^, and sequencing the entirety of *BRCA1*^34^.

With the release of ONT’s PromethION device, the generation of high coverage, Nanopore clinical WGS data has become much more straight-forward and cost-effective. In this study, we aimed to evaluate the extent to which high coverage, long-read Nanopore sequencing enables the genome-wide analysis of a broad range of somatic variation, by comparison with the current gold-standard, Illumina short-read WGS. We do this through conducting in-depth analysis of germline and tumour sequencing data from a patient with Diffuse Large B-cell Lymphoma (DLBCL), an aggressive form of non-Hodgkin’s Lymphoma. This cancer genome was chosen for detailed study due to the frequent and clinically relevant co-occurrence of both somatic hypermutations and structural rearrangements within DLBCL tumours^35^. DLBCL is also characterised by significant interpatient heterogeneity making the design of targeted sequencing approaches challenging. We find that, while currently available tools are not able to provide reliable genome-wide somatic small variant calls from relatively low-depth Nanopore data, advantages in terms of calling large somatic SVs and phasing multiple mutations are already evident. Additionally, we compare the performance of multiple tools on the long-read data and provide recommendations for future studies.

## Results

The DLBCL patient was recruited as part of a large-scale clinical sequencing study utilizing the Illumina short read platform, details of which have been previously published^10^. This patient was selected for subsequent Nanopore sequencing on the basis that this tumour type has long been recognized to harbor chromosomal abnormalities, such as loss of chromosomal 17p, which can influence clinical prognosis and management. Furthermore, multiple complex rearrangements have been reported in more recent WGS analyses of this tumour type (eg^35^). Peripheral blood and fresh-frozen tumour tissue samples were collected for extraction of germline and tumour DNA respectively (see Methods for details). DNA samples were sequenced using both Illumina HiSeq 150bp paired-end sequencing and long read sequencing on Oxford Nanopore Technologies’ PromethION device. All read sets were mapped to the GRCh37 build of the human genome (see Methods for details). This resulted in ~25X and ~60X coverage of the germline sample on the Illumina and Nanopore platforms respectively, and > 80X coverage of the tumour sample on both platforms. Nanopore reads had a mean length of 4.5kb for the germline sample and 5.2kb for the tumour sample. Further sequencing output statistics are shown in Table 1.

**Table 1:** Summary of sequencing data

| Property | Germline<br>(Illumina) | Tumour<br>(Illumina) | Germline<br>(Nanopore) | Tumour<br>(Nanopore) |
| --- | --- | --- | --- | --- |
| Total bases (Gb) <sup>a</sup> | 90 | 309 | 233 | 348 |
| Total reads <sup>a</sup> | 600,078,362 | 2,066,990,226 | 59,661,315 | 76,741,475 |
| % Bases >Q30 | 87.9 | 88.2 | n/a | n/a |
| % Pass, trimmed<br>bases <sup>b</sup> | n/a | n/a | 81.7 | 74.1 |
| Bases mapped<br>(Gb) <sup>c</sup> | 83 | 284 | 187 | 252 |
| Mean read length /<br>insert size (bp) <sup>c</sup> | 150 / 528 | 150 / 520 | 4,472 / n/a | 5,239 / n/a |
| Error rate (%) <sup>d</sup> | 1.6 | 1.5 | 13.8 | 14.6 |

a For Illumina, numbers given are for total number of pass-filter reads/bases
b For Nanopore, 'pass' reads are those given qscore > 7 by the basecaller. Trimming refers to adapter trimming by porechop.
c For Illumina, these numbers are taken from the deduplicated bam files.
d Mismatches / bases mapped

### Single Nucleotide Variants

Somatic small variant calls from the Illumina (hereafter, short-read) data were generated by running Strelka (v2.0.14.1)^36^ on the tumour and germline samples. There were 7709 somatic variants detected, of which 241 were indels and the rest were single nucleotide variants (SNVs). 2446 of the somatic variants were located within genes. We filtered the short-read somatic small variant calls for clinical relevance using a number of different sources, including COSMIC Cancer Genes Census (v71)^37^, “My Cancer Genome” (www.mycancergenome.org) and ClinicalTrials.gov (http://clinicaltrials.gov), as previously described^10^. This resulted in 13 SNVs, 3 of which were located on chr17 (Supplementary Table 1). These were missense variants in *TP53* and *TAF15*, and a *CD79B* splice donor variant (c.552+1G>A) which RNAseq using MinION showed results in intron retention (see Methods and Supplementary Figure 1).

For the Nanopore (hereafter, long-read) data, we initially ran SNV calling using FreeBayes^38^, as described in^39^, on reads aligning to reference chromosome 17. This chromosome was chosen because it was known to harbour mutations of interest, as described above. FreeBayes was run on the tumour and germline samples separately. After initial filtering to exclude very low quality calls (see Methods), germline calls were subtracted from the tumour calls to obtain a list of 18079 putative somatic SNVs on chromosome 17. More stringent quality filters were then applied (see Methods), leaving 9952 ‘pass’ calls. By comparison, there were 123 somatic SNV calls within chromosome 17 from the short-read data, and only 40 calls were present in both call sets (Figure 1A). While the short-read calls are not expected to exactly reflect the ground truth, this still suggests low sensitivity and very low specificity of calling somatic SNVs in the long-read data. Upon further investigation, we found that 6332 (>60%) of the long-read-only somatic calls were called as germline variants in the short-read data. Hence, the low sensitivity of long-read calling on the germline data resulted in many germline variants being called as ‘somatic’ using this method. To investigate further the poor sensitivity of calling somatic SNVs, we first looked to see how many of the 123 somatic SNV calls from the short-read data were present at each step in the subtraction and filtering of the long-read calls. This revealed that 32/123 somatic SNVs were never called by FreeBayes, while 50/123 had been removed by the initial filtering of the tumour vcf and hence had been called with very low quality in the long-read data. None of the 123 somatic calls had been wrongly removed by germline subtraction and only 1 had been filtered out by the more stringent quality filters post-subtraction. We then compared the properties of the 40 sites called as somatic SNVs by both methods with those of the 83 sites called only with the short-read data. Both sets had very similar median values of depth, average base quality and average mapping quality (Figure 1B), hence these properties could not account for the poor sensitivity of somatic SNV calling in the long-read data. Encouragingly, all 3 of the somatic SNVs of clinical interest on chromosome 17 were included in the calls detected by both methods.

**Figure 1.**
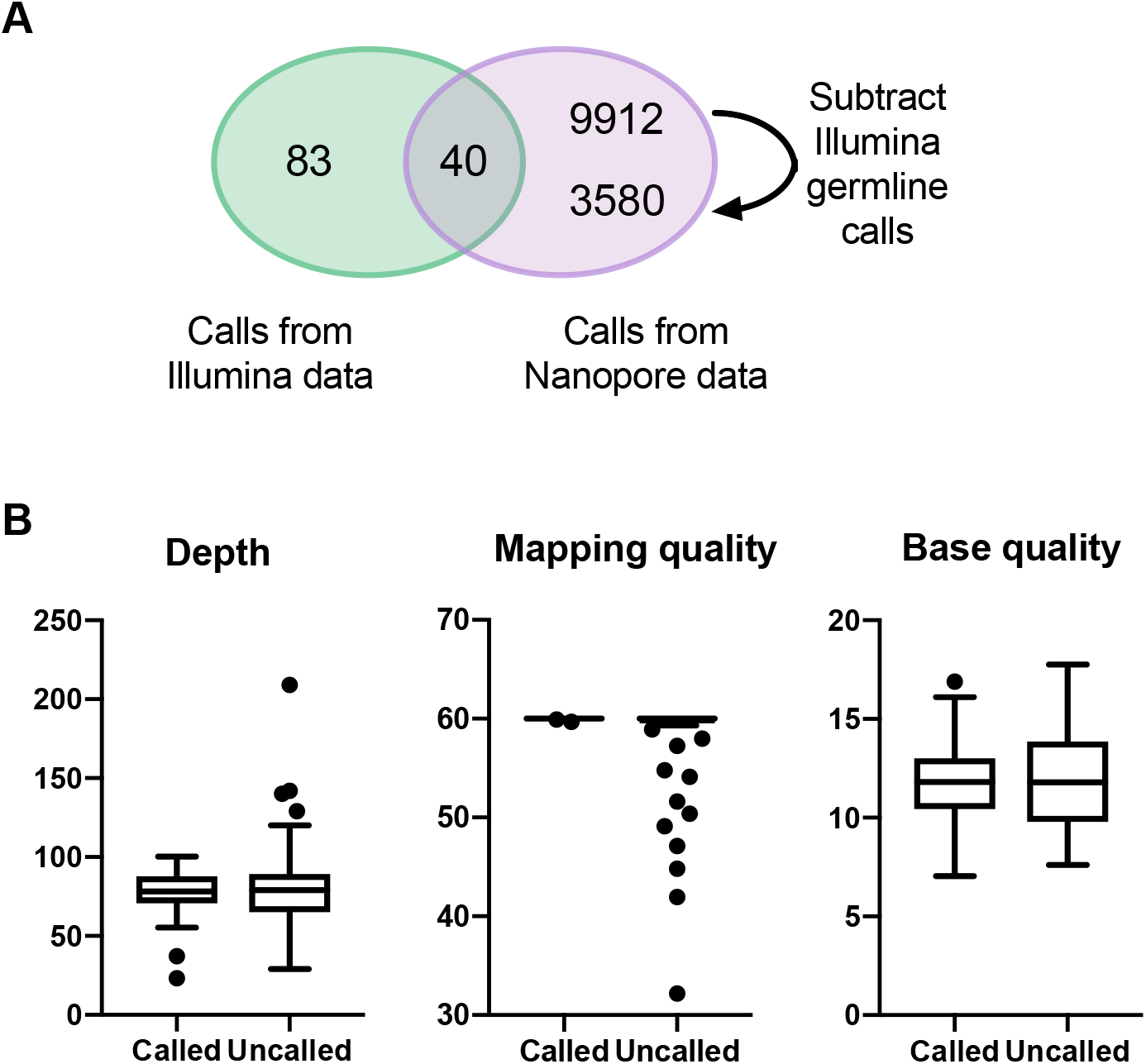
Comparison of somatic SNV calls from short-read and Nanopore data sets. A: Venn diagram to show the overlap between somatic SNV calls from the short-read (green) and Nanopore (purple) data for chromosome 17. The two numbers in the Nanopore-only sector show the number of calls before (top) and after (bottom) subtraction of short-read germline calls. B: Read depth, average mapping quality and average base quality calculated from Nanopore reads covering each of the 123 sites at which an SNV is called in the shortread data. Sites are portioned into those at which an SNV was called vs uncalled in the Nanopore data. Boxes span the interquartile range, with the median marked as a horizontal bar, while whiskers mark the farthest points that are not outliers.

To check that these results were not particular to chromosome 17, we then applied the same methods to reads mapping to chromosome 22, obtaining very similar results in terms of the amount of overlap between short and long-read calls (see Supplementary Figure 2A). Additionally, we repeated the analysis of chromosome 17 SNVs using a second variant caller, Clairvoyante^40^. Clairvoyante is a recently pubished neural network-based variant caller which includes a model trained on long-read data. We used this model to generate germline and tumour SNV calls separately as before, subtracting the germline calls from the tumour calls and filtering the resulting somatic calls based on QUAL scores. Results, in terms of indicated levels of sensitivity and specificity, were similar to those obtained with FreeBayes. For example, using a QUAL score cut-off of 180 resulted in 9987 putative somatic SNVs, only 40 of which overlapped with the short-read calls (full results shown in Supplementary Figure 2B).

### Large structural variants and copy number abnormalities

We compared three structural variant (SV) callsets; calls generated from short-read data using Manta^41^, calls generated from long-read data using the long-read SV caller, Sniffles^42^ following mapping with minimap2^43^, and calls generated from long-read data using Sniffles following mapping with ngmlr^42^ (see Methods). We focused on large structural variants >10kb, examining each in turn to classify them as either true (genuine somatic abnormalities) or false (germline variation or bioinformatic artefacts). There were 39 deletions, 9 inversions, 5 duplications and 3 translocations detected in both the short-read and long-read data, all of which were assessed to be true (Figure 2A). No large insertions were detected by any method. SV calls of different types were unevenly distributed across the genome. In particular, there was a high density of inversions and duplications located on chromosome 16 (Figure 3).

**Figure 2.**
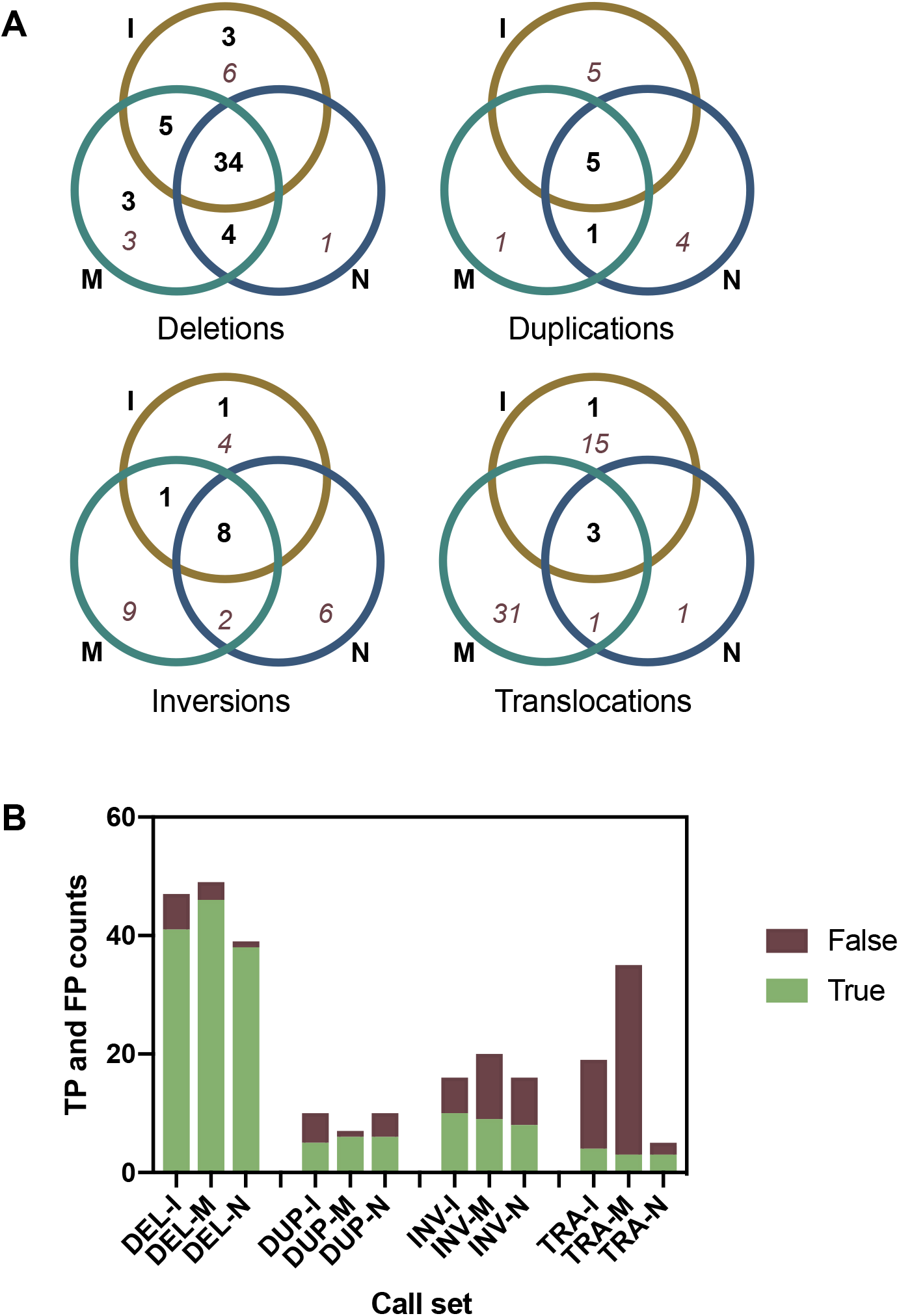
Large (>10kb), somatic structural variant calls obtained via different methods. A: Venn diagrams showing overlap between calls obtained from Illumina short-read data (I; ochre), Nanopore long-read data mapped with minimap2 (M; turquoise) and Nanopore long-read data mapped with ngmlr (N; blue). Bold, black numbers represent true positive calls, while italic, dark red numbers represent false positive calls. B: Data as above for (A), but shown as total counts for each method in a bar chart. Calls are grouped into deletions (DEL), duplications (DUP), inversions (INV) and translocations (TRA). True positive (TP) calls are shown in green and false positive (FP) calls are shown in red-brown.

**Figure 3.**
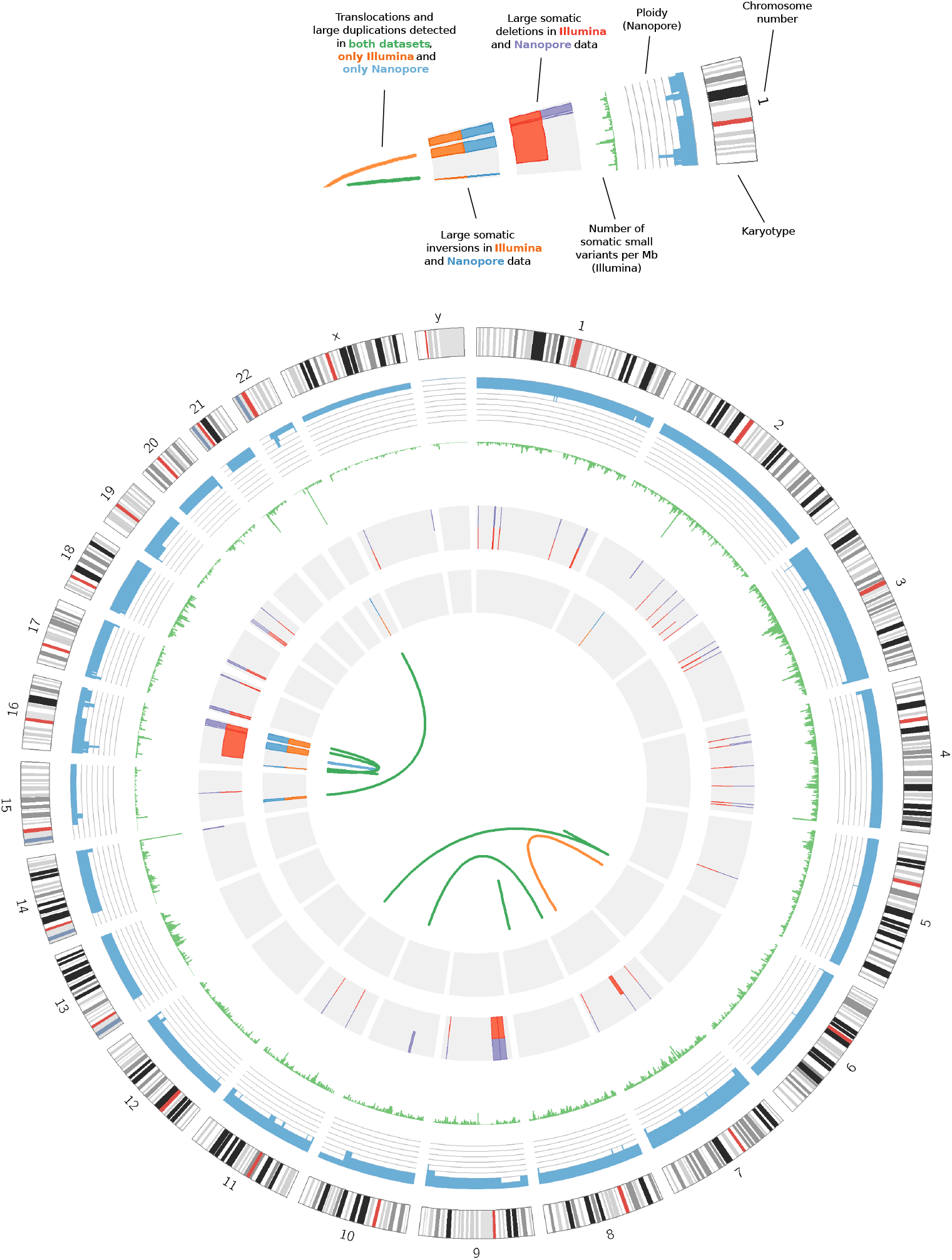
Circos^44^ plot. Rings from outside in: (1) Human karyotype for assembly GRCh37. (2) Ploidy of the tumour sample (blue), with radial axis ranging from 0-8 and grey lines corresponding to each integer value. Data generated from Nanopore read counts (see Methods). (3) Number of somatic small variants per Mb in the Illumina short-read data (green). (4) Large somatic deletions detected in the Nanopore data (purple, outer half) and Illumina short-read data (red, inner half). Only those called assessed as true somatic variants are shown. (5) Large somatic inversions detected in the Nanopore data (blue, outer half) and Illumina short-read data (orange, inner half). Again, only true somatic variants are shown. (6) Links denoting verified translocations and duplications detected in both datasets (green), one translocation detected only in the Illumina short-read data (orange), and one duplication detected only in the Nanopore data (blue)

Across the whole genome, there were 3 deletions, 1 inversion and 1 translocation that were called only in the short-read data but assessed to be true somatic variants (Figure 2A). Upon closer examination of the long-read alignments in IGV, all these SVs also had read support in the long-read data, but at too low a frequency to be called. In two cases this was down to low coverage (defined as < = 10 reads) at one of the breakpoints.

Conversely, 7 deletions and 1 duplication were detected in the long-read data but not the short-read data and visually assessed as being real (Figure 2A). These calls had been missed in the short-read data for a variety of reasons including low allele fraction, not passing the somatic score threshold, the presence of other small SVs nearby, and breakpoints being located in a segmental duplication resulting in miscalling (example given in Supplementary Figure 3).

We estimated false discovery rates (FDRs) and false negative rates (FNRs) for all SV calling methods, based on the assumption that all large somatic SVs in this sample were detected by at least one of the methods. The lowest FDRs and FNRs were achieved using the long-read data for deletions and duplications, and using the short-read data for inversions and translocations. There were noticeable differences between the SV calls resulting from the two long-read mappers, but these depended on the category of SVs being considered. Considering all SV types together, lower FDRs but higher FNRs were achieved when using ngmlr (Figure 2A & B; Table 2). The differences in false discovery rate were most evident in the translocations category, while the differences in false negative rate were most evident for the deletions. In other words, we found that mapping with ngmlr lead to much better specificity for calling somatic translocations, while mapping with minimap2 led to much better sensitivity for calling somatic deletions.

**Table 2:** Estimates of false negative rates (FNR) and false discovery rates (FDR) for different structural variant calling methods. Bold numbers highlight the lowest values in each column

| Method | Deletions |  | Duplications |  | Inversions |  | Translocations |  | All |  |
| --- | --- | --- | --- | --- | --- | --- | --- | --- | --- | --- |
|  | FNR | FDR | FNR | FDR | FNR | FDR | FNR | FDR | FNR | FDR |
| Illumina + Manta | 0.14 | 0.13 | 0.17 | 0.5 | **0** | **0.29** | **0** | 0.79 | 0.12 | 0.32 |
| ONT + Minimap2 + Sniffles | **0.06** | 0.06 | **0** | **0.14** | 0.1 | 0.55 | 0.25 | 0.91 | **0.07** | 0.42 |
| ONT + ngmlr + Sniffles | 0.22 | **0.03** | **0** | 0.4 | 0.2 | 0.5 | 0.25 | **0.4** | 0.20 | **0.21** |

We additionally called copy number abnormalities (CNAs) by calculating the Log2R of read counts in the tumour vs germline sample, in sliding windows across the genome (see Methods for details). This revealed that the ploidy of chromosomes 3, 7, 18 and Y differed in the tumour sample, changing to 4, 3, 3 and 0 respectively (Figure 3). These changes in ploidy were detected in both the short-read and long-read data. There were also three smaller CNAs that were called in both datasets by analysing the Log2R of read counts but had not been detected by Manta/Sniffles. These were a terminal copy number (CN) loss starting at chromosome 10q24.2 and interstitial CN losses in chromosomes 15q13.3-q14 and 22q11.1-q11.21. For the variants on chromosomes 10 and 22, breakpoints corresponded with regions of zero coverage, which explained why they were missed by the SV callers. In the case of the chromosome 15 CN loss, the end breakpoint had been reported as part of an inversion, and the start breakpoint also showed split reads, but with secondary mappings to the decoy chromosome which had not passed filtering in our SV calling pipelines.

We noticed that several of the inversions reported in this data set only had read support for one breakpoint, suggesting they were part of a larger SV complex. An advantage of the long reads is the ability to phase and piece together such complex SVs. To demonstrate this capability, we looked in further detail at chr16:3,213,000-3,894,000, a region containing two large ‘inversions’ with end breakpoints within 300bp of each other, as well as a 6kb deletion. WhatsHap^45^ was run using germline SNP calls from the short-read data to phase the long reads. All reads that could be phased and supported the deletion and inversions were assigned to the same haplotype, confirming that the multiple SVs lie *in cis* (Figure 4A). Hence we were able to reconstruct the highly rearranged somatic haplotype and infer the copy number changes at higher resolution than given by raw CNV calling output (Figure 4B).

**Figure 4.**
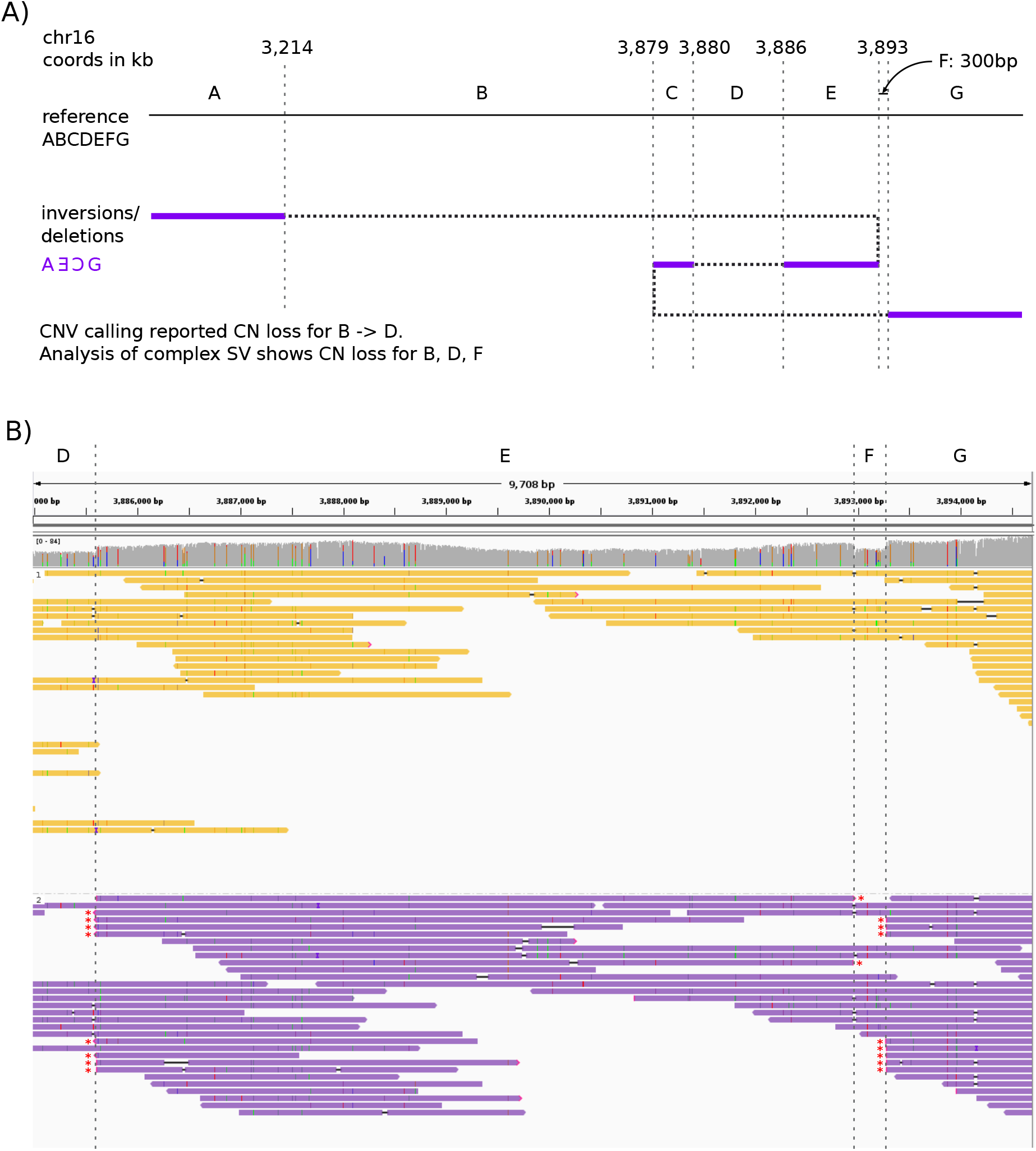
A) Diagram to show the reconstruction of a complex somatic structural rearrangement comprising three deleted segments and two inverted segments. Top: arrangement of segments in the reference sequence, and coordinates of SV breakpoints given to the nearest kilobase. Breakpoints are indicated by vertical dashed lines. Bottom: The rearranged haplotype is obtained by following the path of the dotted/purple line, with purple sections indicating the included segments. The inversion breakpoints between segments A-E and segments C-G were reported by both Manta (short-read data) and Sniffles (long-read data). B) Integrative Genomics Viewer screenshot showing Nanopore reads for the tumour sample, grouped and coloured by haplotype as determined by WhatsHap. Again, vertical dashed lines indicate the position of breakpoints and segments are labelled with letters as in (A). Split ends supporting the breakpoints are indicated with a red asterisk. All reads supporting all 3 breakpoints are assigned to the lower (purple) haplotype, confirming all breakpoints in (A) as in cis. Note that it would not have been possible to confidently phase these breakpoints from the short-read data alone. The nearest heterozygous SNVs to section F are at 3,887,048 and 3,895,908. This means it is not possible to say whether the breakpoints at A|E and E|C are in phase, nor whether A|E and C|G are in phase.

### Large variants of clinical interest

Acquired copy number (CN) events detected in the DLBCL genome ranged in size from ~10kb to ~79.86Mb (Supplementary Table 2). A total of 66 CN events included one or more genes annotated in the cross-referenced cancer-related gene lists. Of particular note were events involving genes reported previously as disrupted in DLBCL. Examples include (i) a heterogeneous, heterozygous CN loss involving *TP53*, (ii) a homozygous loss involving *CDKN2A/2B*, noted previously in 30% cases of activated B-cell like (ABC) DLBCL^46,47^ (iii) a heterogeneous, heterozygous CN Loss involving the acetyl transferase gene *CREBBP*, somatically mutated or deleted in ~25% cases^48,49^ (iv) the more rarely observed CN Loss of *P2RY8* (v) a high CN gain involving the proto-oncogene *BCL6* and *PIK3CA* (vi) a high CN gain involving *CIITA*, mutation of which is implicated in tumour immune escape^50^ (vii) a CN gain involving *IRF4*, involved in terminal differentiation^51^(viii) a CN gain involving *CARD11* in the BCR signaling pathway, mutated in ~9% of ABC-DLBCL cases^52,53^ (ix) a high CN gain involving *MYD88*, encoding an adaptor molecule critical for signal relay from the TLR to the NF-κB transcription complex, as well as the interferon and JAK/STAT3 signaling cascade^54,55^.

On reviewing these findings, the heterogeneous, heterozygous 1.6Mb CN loss encompassing *TP53* was of particular interest because it was found in combination with a somatic missense change in a *TP53* exon (p.Arg249Met; ClinVar accession VCV000376653.2). Clinically, *TP53* mutations or losses have been associated with high grade malignancies and have been found in ~20% of DLBCL patients^56,57^. Therefore, it was of interest to find out if we could determine from our sequencing data whether the point mutation and the deletion were in *cis* or in *trans* (i.e. arising on different alleles or occurring within the remaining undeleted alleles of the chromosome from which the CN loss originated). In the tumour sample as a whole, the deletion was estimated to occur at an allele fraction of 22% in the short-read data and 27% in long-read data. The tumour sample purity was estimated at ~60% (see Methods). Given this level of purity, we would expect a somatic SNV on the deleted chromosome to appear at much less than 30% frequency, whereas a somatic SNV on the non-deleted chromosome could appear at up to 43% frequency, depending on the level of subclonality. The somatic variant appeared at 44% and 52% frequency in the short-read and long-read data respectively, supporting the *in trans* orientation. We wished to add further weight to this conclusion through phasing. However, since the nearest heterozygous germline SNPs were more than 5.5kb away, this was not possible with the short-read data. In the long-read data (which had an average read length of 5.2kb; Table 1), we were able to find three reads containing the somatic variant and spanning to one or other of the heterozygous SNPs. In each case the somatic variant was in phase with what appeared to be the non-deleted alleles (Figure 5A). We were able to extend the phase block further in the 5’ direction by running WhatsHap^45^ on the long-read data (Figure 5B). The allele counts at 13/14 heterozygous germline SNPs within this phase block were suggestive of this haplotype corresponding to the non-deleted chromosome. The counts at the remaining site (rs2908807) were inconclusive likely due to a high frequency of errors. Taken together, evidence from the long-read data gives high confidence that the p.Arg249Met variant and deletion are in *trans*. Confirmation of biallelic mutations in *TP53* may be clinically important for assessing prognostic and therapeutic implications.

**Figure 5.**
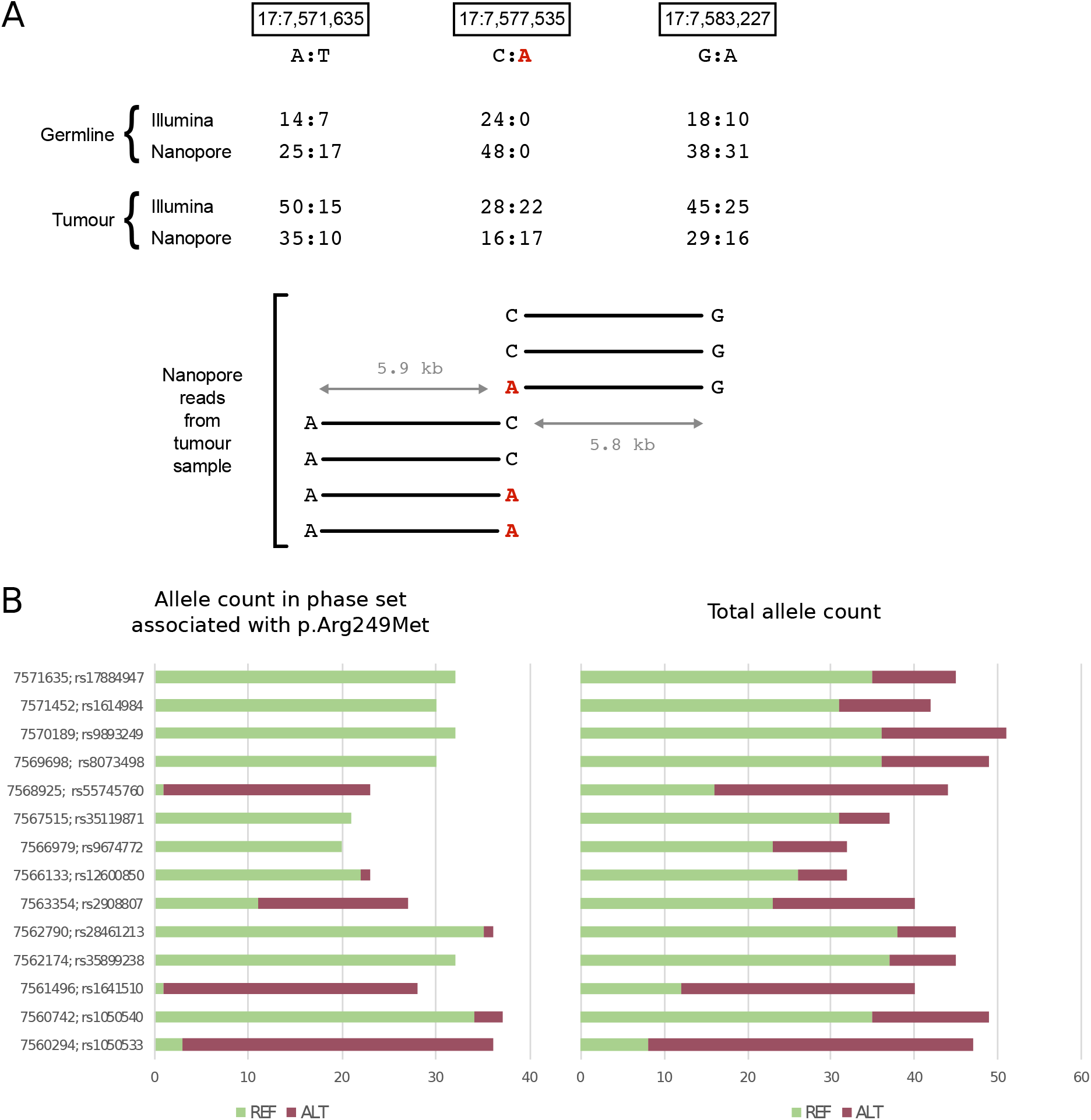
A) Read data used for phasing a somatic variant and large deletion in chromosome 17. Top half: The locations of two heterozygous SNPs (left and right) and somatic single nucleotide variant, p.Arg249Met, (centre) are shown. All three lie within the region covered by the deletion. Base counts are shown reads from both samples run on both platforms (for Nanopore, the reads mapped with minimap2 were used). The somatic variant is highlighted in red. Bottom half: 7 reads spanned from the site of the somatic variant to one or other of the heterozygous SNPs. Each read is shown as a horizontal bar with the base calls at the relevant sites annotated. The approximate distance between the sites is indicated by the grey arrows and text. B) Counts of REF (green) and ALT (dark red) alleles at sites of germline heterozygous SNPs within the range of the phase set covering 17:7,577,535 and in which the somatic mutation p.Arg249Met is seen. Allele counts are given for only reads within the phase set (and assigned to the same haplotype), as given by WhatsHap (LHS), and for all reads covering the position (RHS). The chromosome 17 coordinates and SNP id are given on the extreme left. Although this phase block extends ~17kb in the 5’ direction from the somatic mutation, this does not reach the distal breakpoint of the deletion, which is ~166kb away.

## Discussion

Long-read sequencing is becoming increasingly used in clinical research^58^. Advantages over short-read methods have been noted for applications such as identifying SVs^28,59^, resolving complex SVs^29^, phasing alleles^60^, sequencing repetitive or highly homologous regions^61^ and inferring methylation state^30^. The fast turn-around time of Nanopore sequencing in particular has already led to the development of rapid diagnostic assays for specific cancers^30,31^. However, the question of whether Nanopore long-read sequencing should replace or complement short-read sequencing for cancer diagnosis more generally remains open.

Our work highlights one of the main issues preventing a switch-over to Nanopore long-read sequencing; the difficulty of genome-wide SNV calling with error-prone long-reads alone. In this study, we obtained high coverage long-read data (~ 60X germline and ~80X tumour) of samples from a patient with diffuse large B-cell lymphoma and employed the latest published methods for genome-wide SNV detection. We found that the low sensitivity and specificity of these methods seriously hampers their use for the detection of somatic SNVs, highlighting the need for sophisticated joint calling algorithms that can handle long reads and high error rates. It should be noted that poor genome-wide SNV calling results do not preclude the use of the same long-read data for calling variants (even somatic variants) at specified sites of interest, as shown by the three clinically relevant chromosome 17 somatic mutations that were correctly called in this study.

When it comes to structural variant calling, error rates are less problematic and current long-read tools perform well on Nanopore data. In this study, we examined a total of 158 potential large somatic SVs reported by short- and long-read methods and found the overall sensitivity and specificity of long-read methods for detecting large somatic SVs is similar to that of short-read methods. However, the results vary by SV type, and our analysis highlights the strengths and weaknesses of various tools. From this we can draw a couple of recommendations for those wishing to use Nanopore data to detect somatic SVs: (1) only consider ‘PRECISE’ calls reported by Sniffles, and (2) minimap2 should be used for mapping, except for the detection of translocations (where ngmlr performs better). These recommendations are preliminary and need to be confirmed by the study of additional samples. In examining all large somatic SV calls produced from the short-read data, we found that all those that were “missed” in the Nanopore SV calling appeared clearly supported by the long reads when visually inspected in IGV This would suggest that the information needed for more sensitive and specific calling of structural variants is readily available in long-read data and hence SV calling performance can be expected to improve as tools are fine-tuned based on the availability of more data.

In this study we focused on SVs over 10kb in length, as SVs of this size are more likely to be acquired and not germline SVs. However, others have reported that the largest improvements in sensitivity gained by using long reads for SV calling are seen in the 50-2000bp size range^27,28,62,63^. Individually assessing all these medium-size somatic SV calls for validity is not within the scope of this study, but it is reasonable to expect that with somatic SVs we would also see increased sensitivity compared to short read methods at smaller length scales.

We have provided a couple of examples showing the advantages of long reads for phasing. Firstly, long reads improve our ability to resolve complex structural variants, as shown with the chromosome 16 example (Figure 4). Complex SVs are common in cancer and resolving them can be critical for interpreting pathogenicity^18,29^. Secondly, we also used long read information to phase a somatic mutation (p.Arg249Met in *TP53)* and large deletion encompassing the same gene, confirming that they were *in trans*. Although it is not clearly established whether bi-allelic *TP53* mutations have a worse prognosis compared to monoallelic disruption, this seems logical as TP53 forms homo-dimers and expression is bi-allelic. Deletion of one allele will therefore lead to decreased expression of a normal TP53 protein combined with expression of an abnormal protein from the other allele. In both of these cases high-quality germline SNV calls (from short-read data) were used to extend phase blocks with WhatsHap, but simply examining long reads spanning the variants of interest visually would have been sufficient for giving good phasing confidence.

ONT devices, firmware and tools are continually being updated, and consequently most results may no longer reflect the state-of-the-art by the time they are published. The Nanopore sequencing of these samples commenced at the beginning of 2018, and a PCR-based workflow was chosen in order to achieve high coverage with the available number of flowcells. Since that time, we have seen more than 10-fold increases in yield on the PromethION, and now routinely obtain 30X coverage of the human genome (90Gb mapped data) from a single flowcell of PCR-free library, with mean read lengths of 10-12kb. We have also seen a decrease in mean error rate from 14% to 10%, following firmware and basecalling updates. ONT have recently released the R10 version of the flowcells which are anticipated to lead to additional improvements in the sensitivity and specificity of SNV calling, getting closer to that obtained with short-read data. It is worth pointing out that deep learning tools such as Clairvoyante may not necessarily achieve better results given higher quality input data until they have been re-trained on similar higher quality data. Secondly, we would expect more uniform coverage of the genome, with fewer ‘dark’ regions^26^ as a result of longer read lengths and the removal of the need for PCR amplification. This would enable variant calling within these previously ‘dark’ regions, which contain a substantial number of disease-relevant genes, and would allow us to phase across longer regions.

Nanopore sequencing is not the only long-read technology available. Pacific Biosciences have recently launched a high-fidelity (HiFi) long-read sequencing approach which can achieve mean per-read accuracy of 99.8%^64^. These long read data are likely to hugely improve SNV calling results, however the method requires high input amounts of DNA (e.g. >15ug for 11kb insert sizes^65^) which would be a major drawback for some clinical applications. Both of these technologies are exciting and have potential to expand the horizons of clinical research, however the unstable nature of new technologies represents a challenge for the benchmarking of sequencing results and downstream analysis tools. Additionally, costs, while decreasing, remain high for now; generating 30X coverage of a genome with long-reads is still several times more expensive than with short-reads. Even considering these issues, the power of Nanopopre sequencing to detect structural variants has already led to it being used in large scale population sequencing projects^66^. Our results suggest it as a useful complementary approach for cancer genome sequencing also.

## Methods

### Patients and Ethics

The patient was consented as part of a local clinical WGS programme for analysis of tumor and constitutional DNA (as previously described in^10^). Feedback of clinically actionable germline variants was optional. Written informed consent was obtained in line with the Declaration of Helsinki and local research ethics committees following the procedures outlined by the Oxford Radcliffe Biobank (South Central-Berkshire B Research Ethics Committee [REC no: 14/SC/1165]).

### Sample Preparation and DNA Extraction

#### Tumor Tissue Handling

The patient underwent biopsy of primary and/or metastatic cancer to obtain fresh tissue for sequencing. Briefly, fresh tissue samples were collected at the time of resection of the abdominal lymphomatous mass. Samples were snap-frozen in liquid nitrogen and an H&E frozen histology section was taken to confirm tissue content. Only samples with microscopically estimated tumor cell content of >40% were used for sequencing. Frozen tissue was thawed rapidly for nucleic acid extraction and sequencing for Illumina following procedures previously delineated in^10^.

#### Long-read library preparation: tumour DNA

2 μg of lymphoma DNA was thawed fragmented in 49 μl of Nuclease Free Water (NFW) for 2 minutes at 7,000 revolutions per minute (rpm) using g-TUBE (Covaris^®^, Woburn, MA, USA) following manufacturer recommendations. FFPE repair, End repair-dA tailing and PCR Adapter (PCA) ligation, were performed following Oxford Nanopore Technologies’ (ONT) 1D Low input genomic DNA with PCR (SQK-LSK108) version LIP_9021_v108_revL_11Nov2016. End repair incubation times were extended to 30 minutes at 20°C and 30 minutes at 65°C. Incubation of DNA with Agencourt AMPure XP beads (Backman Coulter Inc., Brea, CA, USA) and elution times were increased to 20 and 10 minutes at room temperature respectively. Pre-PCR size selection was performed as follows: After washing the beads with 70% ethanol, the PCA ligated DNA was re-suspended in 100 μl of NFW and transferred to a new 1.5 μl Lobind Tube (Eppendorf, Hamburg, Germany). Avoiding pelleting the beads on magnetic bar, an additional volume of 40 μl of beads was added to the previous re-suspended material resulting into a 0.4X ratio of beads to DNA volume. The tube was incubated on the Hula Mixer (Thermo Fisher Scientific, Waltham, MA, USA) for 20 minutes at room temperature. The beads were washed twice with 200 μl of 70% ethanol and re-suspended in 28 μl of NFW followed by 10 minutes elution at room temperature. The PCA ligated and size selected DNA was diluted to a concentration of 10 ng/μl in NFW. The PCR reaction was set up as follows: 46 μl Nuclease-free water, 2 μl Primers (PRM, Oxford Nanopore Technologies, Oxford, UK), 2 μl 10ng / μl adapter ligated template, 50 μl LongAmp Taq 2x master mix (New England Biolabs, Ipswich, MA, USA). Initial denaturation was for 3 min at 95 °C, denaturation 15 secs at 95 °C (15 cycles), annealing 15 secs at 62 °C (15 cycles), extension 15 min at 65 °C (15 cycles), final extension 15 min at 65 °C, hold at 4 °C. PCR products were cleaned up with a 0.4X ratio of beads to DNA volume and eluted in 25 μl NFW. Five aliquots of 1.5-2 μg size selected and amplified tumour DNA were prepared and sequenced according to the ONT protocol kit 9 chemistry version GDLE_9056_v109_revC_02Feb2018. These were run on four PromethION flow cells using the MinKNOW software version 1.18.02 for 64 hours.

#### Germline DNA Extraction

Germline DNA was isolated from 1.5 ml peripheral blood using the QIASymphony DSP DNA Midi kit (QIAGEN), according to the manufacturer’s protocol. Illumina short-read libraries of 350-bp fragments were generated and sequenced as described in^10^. 2 μg of germline DNA from the same lymphoma patient were treated following the same pre-PCR procedures as per the tumour sample above. Post-PCR procedures were completed according to a newly updated version of the ONT SQK-LSK109 kit protocol, GDE_9063_v109_revC_04Jun2018. Five 1-1.5 μg of size selected and amplified germline DNA aliquots were run on four PromethION flow cells using ONT MinKNOW software version 1.18.02 for 64 hours.

In parallel, two additional R9.4.1 MinION flow cells were run using 1 μg of size selected and amplified germline and tumour DNA following the SQK-LSK109 protocol version GDE_9063_v109_revA_23May2018 (Oxford Nanopore Technologies, Oxford, UK).

#### RNA lymphoma cDNA library preparation

The lymphoma transcriptome was processed according to the Oxford Nanopore SQK-PCS108 cDNA-PCR sequencing protocol (version PCS_9035_v108_revD_26Jun2017, last update 25/10/2017). We used 500ng of total RNA into a final volume of 9 μl. Note that due to the low input of the original clinical aliquot, we could not recover the optimal starting input of ~50ng of PolyA RNA. The mRNA was targeted for reverse transcription using Nanopore oligo-dTs. Subsequently, full-length double stranded cDNA was generated employing the Nanopore Strand Switching Primer and amplified in 50 μl reaction volumes.

The PCR reaction contained 5 μl cDNA, 25 μl 2X LongAmp^®^ Taq Master Mix (NEB Biolabs^®^ Inc, Ipswich, Massachusetts, USA, cat. #M0287S), 1.5 μl Nanopore cDNA Primers and 18.5 μl NFW. The reactions were incubated at 95 °C for 30 seconds, followed by 15 cycles of (95 °C for 15 seconds, 62 °C for 15 seconds, 65 °C for 8 minutes) with a final extension at 65 °C for 6 mins. The amplification products were normalised to ~400 fmol into 20 μl of Nanopore Rapid Annealing Buffer. The cDNA was adapter ligated adding 5 μl of Nanopore cDNA Adapter Mix for 5 minutes at room temperature. The adapter ligated library was subsequently cleaned-up by adding 40 μl AMPure XP beads, incubating for 5 minutes at room temperature and re-suspending the pellet twice in 140 μl of Nanopore Adapter Binding Beads. The purified ligated cDNA was then eluted in 12 μl of Nanopore Elution Buffer. The library was run on a MinION R9.4.1 flow cell using the 48 hour PCS108 MinION sequencing script.

### Bioinformatic analysis of the Illumina data

Analysis was performed using a bespoke, locked-down, and version-controlled bioinformatics pipeline according to the required specification for clinically accredited laboratories.

Paired-end alignment of sequencing data against the reference genome hg19 (GRCh 37) was performed using the Whole-Genome Sequencing Application v2.0, based on Isaac Alignment Tool, within BaseSpace (Illumina). Somatic single nucleotide (SNV) and insertion/deletion (InDel) variant calling analysis was performed using the Tumour-Normal Application v2.0, based on Strelka, within BaseSpace. Calls were annotated using variant effect predictor (VEP) within Ensembl-tools v84, including COSMIC v71.

Manta (v0.23.1, as part of Illumina’s Tumor Normal Application v2.0.0) was run for somatic structural variant calling.

#### Detection of acquired CN events

WGS bam files (hg19) were analysed for copy number (CN) events using ngCGH (https://github.com/seandavi/ngCGH) for the paired tumour:germline analysis and ngbin (BioDiscovery, Inc., El Segundo, California, USA, available from http://www.biodiscovery.com) for single sample analyses. A window size of 300 reads was applied for ngCGH, and a window size of 1kb and read depth of ten for ngbin. B-allele frequency information was obtained from vcf files generated using Platypus^67^. Singleton and paired tumour-germline Log2R outputs were visualised together with B-allele frequency data using Nexus Discovery Edition 10.1 (BioDiscovery, Inc., El Segundo, California, USA). CN events and regions of homozygosity (ROHs) were flagged using the SNP Rank Segmentation setting. All putative events were visualised and curated manually prior to filtering in Microsoft Excel to remove germline copy number variants (CNVs), germline ROHs and calls due to underlying sequence complexity (eg. segmental duplications, tandem repeat regions) or insufficient confidence on visualisation. For acquired ROHs, referred to as copy neutral loss of homozygosity (cnLOH), the reporting size threshold was >2Mb. All acquired copy number events were cross-checked using IGV, looking for supporting evidence from read pairs mapping to the breakpoint regions.. The final Microsoft Excel file included all filtered events intersected with both database^37^ and ‘in house’ gene lists comprising known genes of interest in cancer

### Bioinformatic analysis of the Nanopore data

Nanopore reads were basecalled with Albacore v 2.3.1 or, for later runs, guppy v 1.6.0 (Oxford Nanopore Technologies; note guppy 1.6.0 contained a similar neural network model to Albacore 2.3.1 but was optimized to run on GPUs). They were then trimmed with Porechop v 0.2.3^68^ and mapped using minimap2 v 2.10^43^. Trimmed reads were additionally mapped with ngmlr v 0.2.7, since the developers of this mapper had reported superior results for structural variant calling^42^.

#### Somatic SNV calling

Reads mapping to chromosome 17 (with minimap2) were selected and split into mapped sections of max 100bp in length. FreeBayes v 1.0.2^38^ was run on the split bams, for tumour and germline samples separately, using the contamination parameters RH/RA=0.7/ 0.1. These parameters had produced good results for variant calling without prior phasing in previous work^39^. Indels and MNPs were not called. Both tumour and germline bams were filtered based on the following criteria: QUAL > 1 & NUMALT < 2 & SAF > 1 & SAR > 1. After this initial filtering, germline calls were subtracted from tumour calls if they shared the CHROM, POS and ALT fields in the vcf. Having previously observed that strand bias is often an indicator of a repeated Nanopore error, we removed calls where the fraction of alternate allele observations was more than 4x higher or lower in the forward strand vs the reverse strand. The resulting somatic vcf was examined for overlap with the short-read SNV calls and the distributions of the following properties were plotted: AO (alternate allele observations), RO (reference allele observations), QA (alternate allele quality sum in phred), QR (reference allele quality sum in phred), MQM (mean mapping quality of observed alternate alleles), AB (the ratio of reads showing the reference allele to all reads), SAP (strand balance probability for the alternate allele), as shown in Supplementary Figure 4. Based on these distributions the following criteria were used for final filtering: AO > 10 & RO > 10 & QA > 100 & QR > 100 & AO < 100 & RO < 100 & SAP < 30 & MQM > 55. The filtering criteria decided on here were applied to the chromosome 22 data.

#### Somatic SV calling

Sniffles v1.0.9^42^ was used to call structural variants with the following parameters: −1 100 -s 10 --enotype. Germline calls were subtracted from tumour calls based on matching ALT fields (i.e. SV type), and both the starts and ends being within (germline SV length)/100 bases of each other. Remaining somatic calls were then filtered according to SVLEN > 10000 & !(CHR2 = hs37d5) & PRECISE. SV calling was done for reads mapped with minimap2 and reads mapped with ngmlr. We examined all long-read and short-read SV calls visually in the Integrative Genomics Viewer (IGV)^69^, with tracks of long-read data, short-read data, and segmental duplications simultaneously loaded. In this way each call was classfied as either true or false depending on whether it was deemed to be a genuine somatic abnormality, or a germline variant or mapping artefact.

#### Comparison of short-read and long-read SV calls

Lists of calls from the three different methods (Illumina + Manta, Nanopore + minimap2 + Sniffles, Nanopore + ngmlr + Sniffles) were examined to decide where the same SV was reported by multiple methods (see Supplementary Table 2). The breakpoints of each unique call were looked at in IGV to assess whether there was sufficient evidence to class a call as a genuine somatic variant. If there was evidence of reads supporting the breakpoint in the germline sample, the call was classed as ‘false’. False discovery rates and false negative rates we calculated by assuming that all true variants were picked up by one of the three methods, and then using the following formulae (TP=true positive; FP=false positive):

FNR = (Number of TPs not detected with this method)/(Total TPs)

FDR = (Number of FPs detected with this method)/(Total calls from this method)

#### Detection of acquired CN events

Aligned reads were split into aligned sections of max 100bp in length, as described for SNV calling above. The number of reads with MAPQ >= 20 within 100kb windows was counted, for both the tumour and germline bam files. Let N(w) be the read count in the germline sample for window, w, and T be defined similarly for the tumour sample. The normalised log read count ratio, R, was calculated as

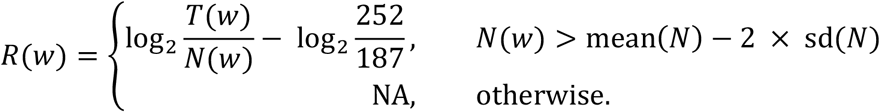

The R package DNAcopy^70^, was used to perform circular binary segmentation and detect change points using the following parameters: alpha=0.01, min.width=5, undo.splits-’sdundo”, undo.SD=3. The mean value of points in all segments longer than 500 windows and representing ploidy=2 (i.e. clustered around 0) was calculated as 0.094. This was subtracted from all segment means, to account for the ploidy of the tumour sample being greater than 2. The purity of the tumour sample was then calculated by similarly computing the mean value of points in segments representing ploidy = 1, 3, 4 (identifiable as distinct clusters of segment means). This gave three separate estimates of purity – 0.61, 0.66 and 0.64, where purity was defined as the amount of tumour DNA in the tumour sample as a proportion of total DNA (tumour + germline). A purity value of 0.63 value was used to calculate the theoretical segment means for values of ploidy from 1 to 6 and then classify the recorded segment means accordingly, for both the autosomes (with ploidy 2 in the germline) and the sex chromosomes (with ploidy 1 in the germline)

#### Phasing

A list of heterozygous SNVs was obtained by running Illumina’s Isaac variant caller on the germline bam. *whatshap phase* was then run using this vcf and Nanopore reads from both the germline and tumour samples. The resulting phased SNV calls were used along with *whatshap haplotag* to assign haplotype and phase set tags to Nanopore reads from the tumour sample. Haplotag bams were visualised in IGV, grouped by haplotype and coloured by phase set.

## Acknowledgements

The Illumina short read sequencing was carried out as part of a larger clinical whole genome sequencing study (OxClinWGS) commissioned by the Health Innovation Challenge Fund (R6-388 / WT 100127), a parallel funding partnership between the Wellcome Trust and the Department of Health. JCT and ATP are funded by the Oxford NIHR Biomedical Research Centre. Computation used the Oxford Biomedical Research Computing (BMRC) facility, a joint development between the Wellcome Centre for Human Genetics and the Big Data Institute supported by Health Data Research UK and the NIHR Oxford Biomedical Research Centre. Financial support was provided by the Wellcome Trust Core Award Grant Number 203141/Z/16/Z. The views expressed in this publication are those of the authors and not necessarily those of the Wellcome Trust or the Department of Health.

## Data availability

The datasets generated and analysed during the current study will be available from EGA, accession EGAS00001004266. These consist of BAM files for Illumina genomes (tumour and normal), nanopore genomes (tumour and normal) and nanopore RNA seq (tumour only).

