## Supplementary Figures and Tables for "Short and long-read genome sequencing methodologies for somatic variant detection; genomic analysis of a patient with diffuse large B-cell lymphoma"

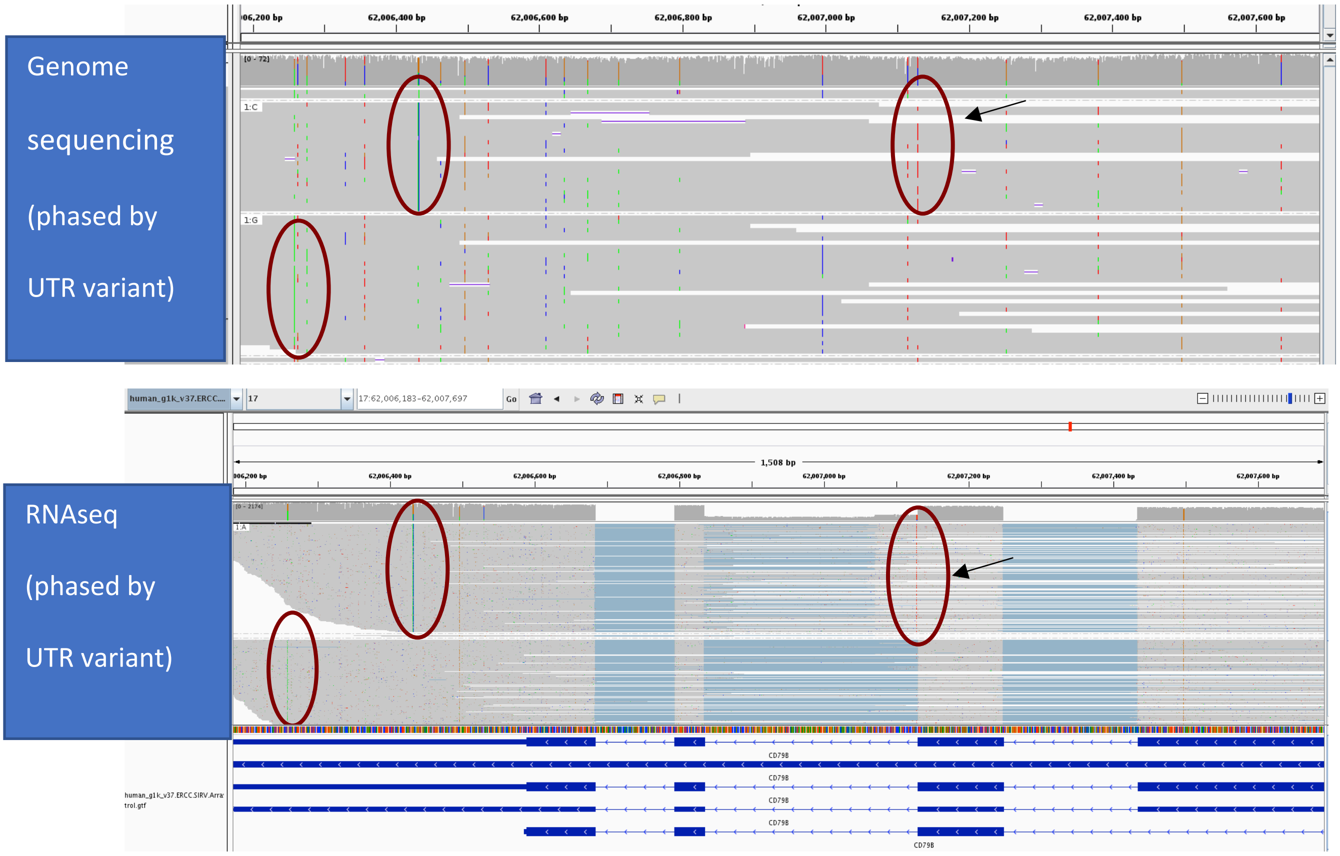


Supplementary Figure 1: The splice donor site mutation (c.552+1G>A, NM 001039933.1) can be accurately phased across multiple exons with the use of long Nanopore cDNA reads. Top: IGV screenshot showing reads from tumour DNA sample, which enable phasing of the somatic variant with one of the 3’-UTR variants. Bottom: Long Nanopore cDNA reads from the tumour RNA sample show that the splice donor variant results in intron retention and partial intron retention. In both screenshots, reads are grouped according to the base at one of the heterozygous SNPs in the 3’-UTR (circled in red). The somatic splice donor site mutation is circled in red and highlighted with an arrow.

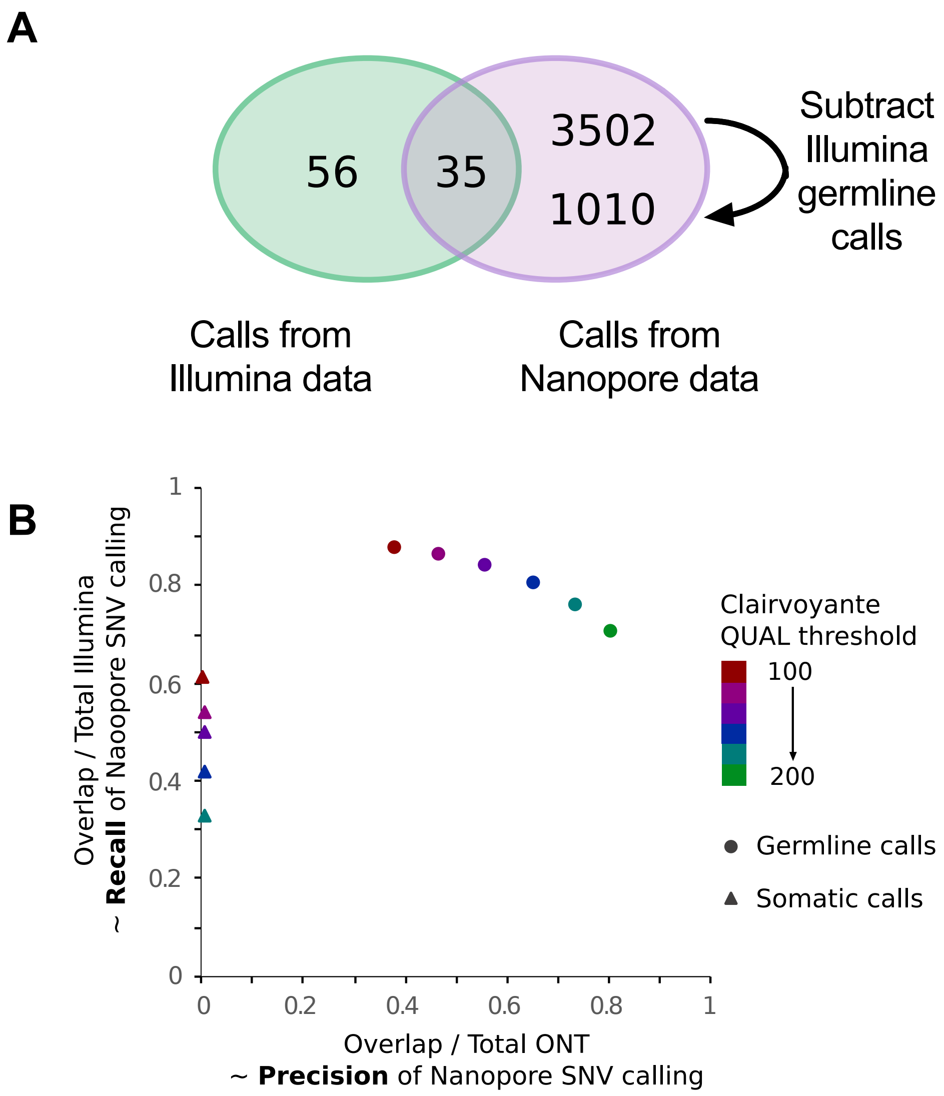


Supplementary Figure 2: A) Overlap between somatic SNV calls generated from Nanopore vs Illumina reads covering chromosome 22. Freebayes was used to call SNVs in the Nanopore data. As with the results for chromosome 17 shown in Figure 1, there is more than an order of magnitude difference between the number of calls in the overlap between the two data sets and the number of calls from the Nanopore data, even after stringent filtering and subtracting the Illumina germline calls. B) The performance of Clairvoyante on the Nanopore data for chromosome 17 is shown. The correspondence between Clairvoyante and Illumina callsets is indicated for germline calls (circles) and somatic calls (triangles). The values displayed on the x and y axes approximate the precision and recall of Nanopore SNV calling respectively, under the assumption that the Illumina calls approximate the truth set. For germline calls the colours represent different QUAL score thresholds used to define the 'PASS' filter (applied uniformly to homozygous and heterozygous calls), from red=100 to green=200. The same colouring is used for somatic calls, but here the variable QUAL score referred to is that used for filtering tumour calls prior to subtraction of Clairvoyante germline calls with minimal filtering applied (QUAL >0). Even following subtraction of this widest set of germline calls a large number of putative somatic calls remained, of which only 0.1-0.4% (QUAL 100 – 180) were found in the Illumina somatic call set.

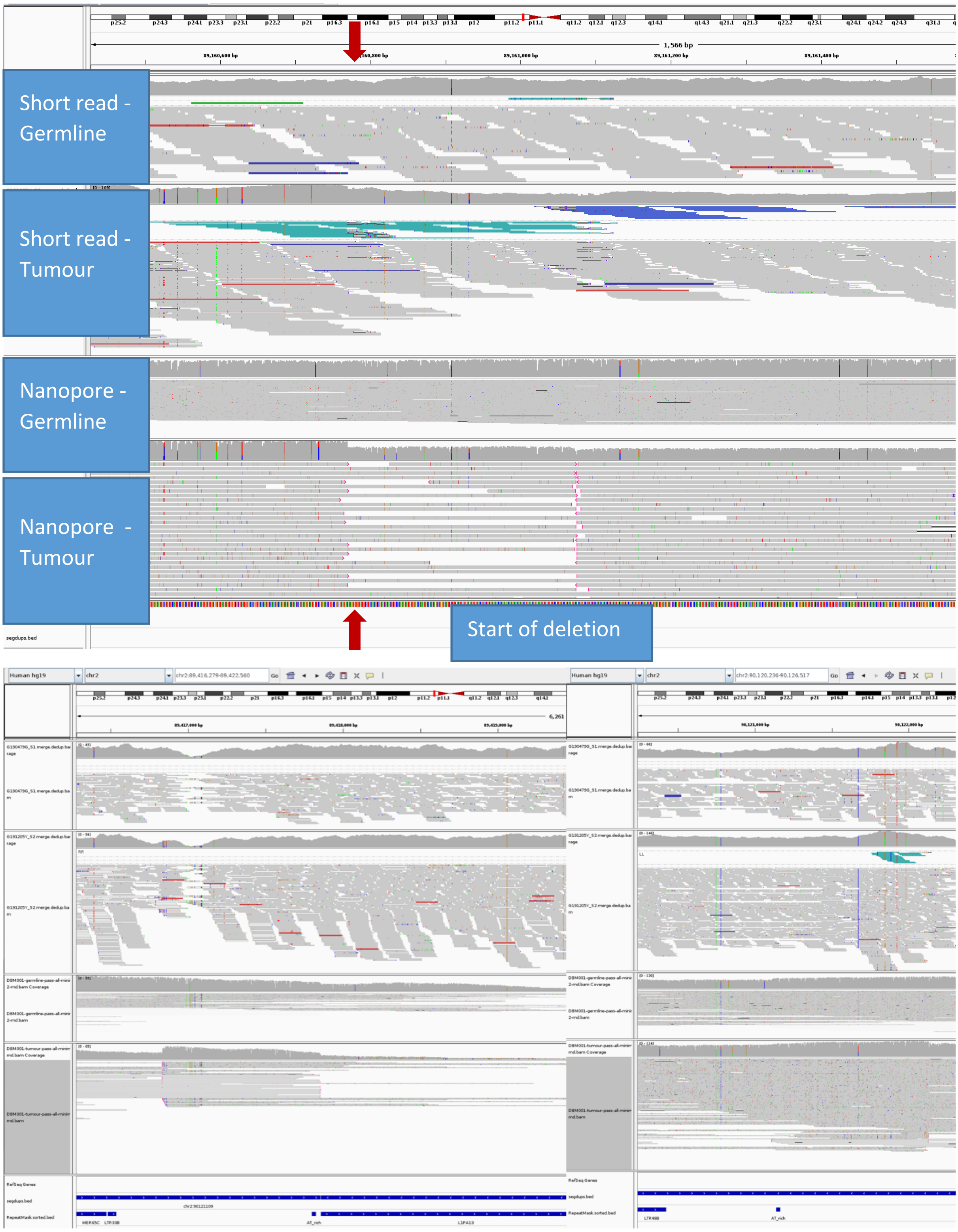


Supplementary Figure 3. Example of a somatic deletion near the centromere on the p arm of chromosome 2, which is incorrectly called as an inversion in the short-read data. Top:

Illumina and Nanopore read alignments at the distal breakpoint (breakpoint of interest indicated by red arrows – note that a second breakpoint corresponding to a smaller inversion can also be seen, and is called as an inversion in both data sets). The four loaded tracks in all screen shots correspond to the Illumina germline bam, Illumina tumour bam, Nanopore germline bam and Nanopore tumour bam, from top to bottom. The long-read alignments clearly show a drop in coverage associated with this breakpoint. Paired-end short reads are coloured according to pair orientation and insert size, hence the many turquoise reads in the Illumina tumour bam suggest an inverted breakpoint, although a few reads (red) support the deletion called in the Nanopore data. Bottom, LHS: IGV screen shot of proximal breakpoint suggested by long-read data; RHS: IGV screen shot of proximal breakpoint suggested by short-read data. The segmental duplications track second from bottom in these screen shots shows that there is a segdup linking these two suggested proximal breakpoints. Several of the long Nanopore reads mapping to the left-hand breakpoint extend beyond the end of the segdup (whereas this is not the case with any of the long reads supporting the right-hand breakpoint), hence suggesting the deletion as the true underlying SV.

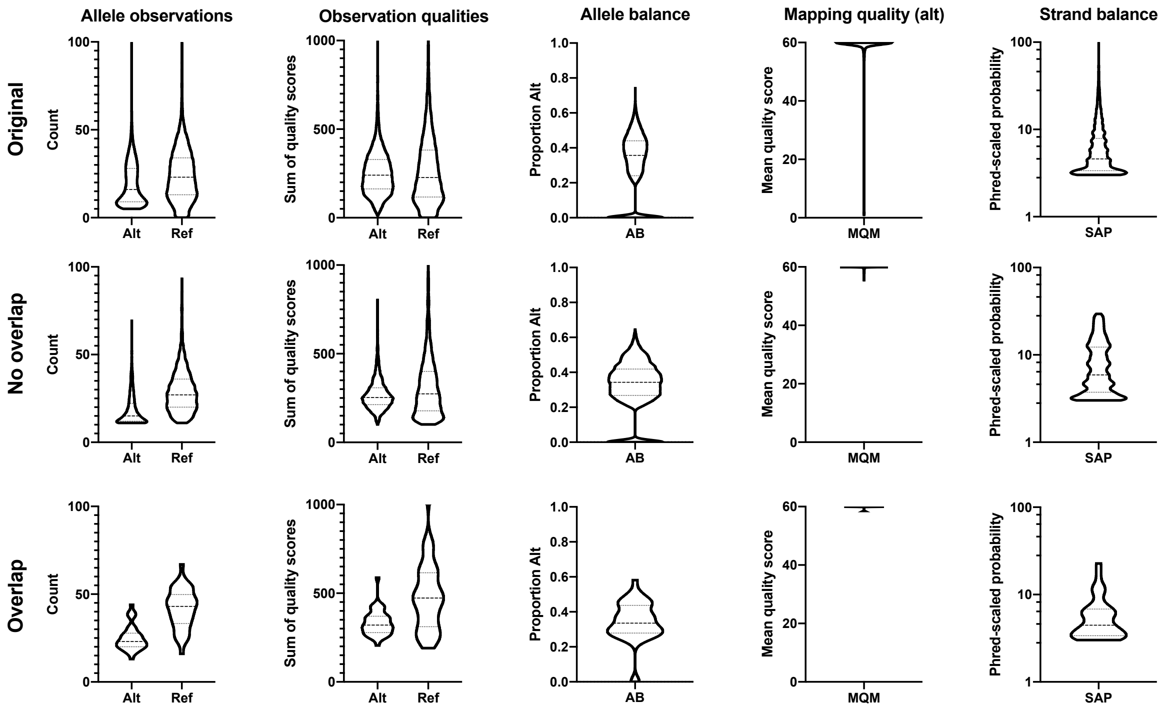


Supplementary Figure 4: Properties of Nanopore SNV calls. Top – all calls. Middle – filtered calls that don't overlap with short-read calls. Bottom – filtered calls that do overlap with short-read calls

**Supplementary Table 1:** Somatic SNVs of potential clinical relevance detected by short-read genome sequencing. Three of the 13 SNVs were located on chr17. Variant annotation was using the Ensembl Variant Effect Predictor web interface (http://grch37.ensembl.org/Homo_sapiens/Tools/VEP), performed on 10th December 2019. Although listed here separately, the three variants in IGLL5 and two variants in IGKV1D-17 lie in close proximity and may represent single mutational events

Supplementary Table 2

| Chr | GRCh37 position | Ref/Alt | Symbol | Transcript ID | Exon (intron) | cDNA annotation | Protein prediction | Existing variation | Pubmed | SIFT | PolyPhen | CADD | MaxEntScan (diff) | gnomAD_AF |
|---|---|---|---|---|---|---|---|---|---|---|---|---|---|---|
| 2 | 90,121,885 | G/C | *IGKV1D-17* | NC_000002.11 | 2/2 | c.103G>C | p.Ala35Pro | - | - | deleterious(0.01) | benign(0.388) | 12.96 | - | - |
| 2 | 90,121,889 | C/G | *IGKV1D-17* | NC_000002.11 | 2/2 | c.107C>G | p.Ser36Cys | - | - | deleterious(0) | probably_damaging(0.987) | 18 | - | - |
| 5 | 256,514 | G/C | *SDHA* | NM_004168.3 | 15/15 | c.1974G>C | p.Pro658 (syn) | rs1042446 | - | - | - | 0.68 | - | 0.0002385 |
| 6 | 37,138,549 | G/A | *PIM1* | NM_001243186.1 | 2/6 | c.356G>A | p.Gly119Asp | rs377274719,COSM1581462 | - | deleterious(0.01) | probably_damaging(0.966) | 23.8 | 1.019 | 3.30E-05 |
| 9 | 139,410,082 | T/C | *NOTCH1* | NM_017617.4 | 11/34 | c.1756A>G | p.Thr586Ala | - | - | tolerated(0.16) | benign(0.006) | 18.02 | - | - |
| 11 | 119,169,084 | C/T | *CBL* | NM_005188.3 | 15/16 | c.2268C>T | p.Ala756 (syn) | rs142564074,COSM923819 | - | - | - | 10.4 | - | 4.38E-05 |
| 14 | 51,224,431 | G/A | *NIN* | NM_020921.3 | 18/31 | c.3317C>T | p.Ser1106Phe | - | - | deleterious(0.02) | possibly_damaging(0.888) | 20.9 | - | - |
| 17 | 7,577,535 | C/A | *TP53* | NM_000546.5 | 7/11 | c.746G>T | p.Arg249Met | rs587782329,COSM129832,COSM129833,COSM1649403,COSM1728798,COSM326723,COSM326724,COSM3388182,COSM3403260,COSM375642,COSM375643,COSM43665,COSM43871,COSM44091,COSM44916,COSM5878478,COSM5878479 | 9569050, 24381225, 24487413, 24641375, 26619011 | deleterious(0) | possibly_damaging(0.868) | 25.7 | - | - |
| 17 | 34,151,181 | G/A | *TAF15* | NM_139215.2 | 7/16 | c.584G>A | p.Arg195Lys | - | - | deleterious_low_confidence(0.02) | benign(0.021) | 23.5 | - | - |
| 17 | 62,007,129 | C/T | *CD79B* | NM_001039933.2 | (4/5) | c.552+1G>A | NA | COSM5045027 | - | - | - | 32 | 8.182 | - |
| 22 | 23,230,312 | C/A | *IGLL5* | XM_005261300.1 | 1/3 | c.79C>A | p.Leu27Met | rs777349722,COSM5946682 | - | tolerated_low_confidence(0.35) | benign(0.117) | 1.081 | - | 1.36E-05 |
| 22 | 23,230,366 | C/T | *IGLL5* | XM_005261300.1 | 1/3 | c.133C>T | p.Pro45Ser | rs575661811,COSM5652743 | - | tolerated_low_confidence(0.21) | benign(0.067) | 1.043 | - | 8.19E-05 |
| 22 | 23,230,381 | C/T | *IGLL5* | XM_005261300.1 | 1/3 | c.148C>T | p.Pro50Ser | rs560170702 | - | tolerated_low_confidence(0.09) | possibly_damaging(0.892) | 7.516 | - | 6.83E-06 |

**Supplementary Table 2:** Somatic SVs detected by one or more pipeline. Breakpoint (BP) positions are given with reference to the GRCh37 genome. SV types include TRA (translocations), DUP (duplications), DEL (deletions) and INV (inversions). Method(s) used to detect SV are indicated by I (Illumina), M (nanopore pipeline with minimap2 mapping) and N (nanopore pipeline with ngmlr read mapping). N/A, SV length not available for translocation events. Comments reflect either the reason a call was missed by some methods (for TRUE calls), or the reason a call could have been falsely made (for FALSE calls), where such reasons were easily classifiable. Background ploidy of the chromosome (of BP1) is included for true calls to aid assessment of whether an SV results in a CN gain or loss (e.g. the deletions in chr 3 are against a background ploidy of 4 and hence result in a CN gain while regions outside the deletions have a high CN gain)

| Chr and position of BP1 | | Chr and position of BP2 | | SV type | Method | Assessment from IGV | SV length | Background ploidy | Comment |
|---|---|---|---|---|---|---|---|---|---|
| 1 | 3,582,983 | 1 | 3,596,979 | DEL | IMN | TRUE | 13,996 | | |
| 1 | 36,587,826 | 1 | 37,928,359 | DEL | MN | TRUE | 1,340,533 | | |
| 1 | 50,242,314 | 1 | 50,265,669 | DEL | IMN | TRUE | 23,355 | 2 | |
| 1 | 171,042,334 | 1 | 171,097,428 | DEL | IMN | TRUE | 55,094 | | |
| 1 | 224,278,927 | 1 | 226,971,073 | DEL | IMN | TRUE | 2,692,146 | | |
| 2 | 89,160,770 | 2 | 89,416,835 | DEL | M | TRUE | 256,065 | | |
| 2 | 89,161,074 | 2 | 89,185,667 | INV | IMN | TRUE | 24,593 | | |
| 2 | 89,203,502 | 2 | 89,544,271 | DEL | MN | TRUE | 340,769 | | |
| 2 | 145,260,038 | 2 | 145,287,029 | DEL | IMN | TRUE | 26,991 | 2 | |
| 2 | 179,801,721 | 2 | 179,938,921 | DEL | IMN | TRUE | 137,200 | | |
| 2 | 201,753,407 | 2 | 201,832,238 | DEL | IMN | TRUE | 78,831 | | |
| 2 | 225,687,648 | 2 | 225,792,718 | DEL | I | TRUE | 105,070 | | Low AF |
| 2 | 242,627,219 | 2 | 242,658,927 | DEL | IMN | TRUE | 31,708 | | |
| 3 | 47,146,368 | 3 | 47,179,903 | DEL | IMN | TRUE | 33,535 | | |
| 3 | 60,454,456 | 3 | 60,775,981 | DEL | IMN | TRUE | 321,525 | 4 | |
| 3 | 61,020,731 | 3 | 61,221,598 | DEL | I | TRUE | 200,867 | | Low Nanopore coverage at 5' end (AT-rich region) |
| 3 | 71,131,337 | 3 | 71,184,317 | DEL | IM | TRUE | 52,980 | | Low AF |
| 4 | 32,894,943 | 4 | 32,943,328 | DEL | IMN | TRUE | 48,385 | | |
| 4 | 46,340,406 | 4 | 46,357,872 | DEL | MN | TRUE | 17,466 | | Small inversion nearby |
| 4 | 48,008,682 | 4 | 48,074,064 | DEL | IM | TRUE | 65,382 | | |
| 4 | 94,754,812 | 4 | 94,792,746 | DEL | IMN | TRUE | 37,934 | 2 | |
| 4 | 136,181,907 | 4 | 136,243,262 | DEL | IMN | TRUE | 61,355 | | |
| 4 | 172,880,509 | 4 | 172,946,865 | DEL | IMN | TRUE | 66,356 | | |
| 4 | 178,262,300 | 4 | 178,275,050 | DEL | IMN | TRUE | 12,750 | | |
| 5 | 117,476,209 | 5 | 117,512,551 | DEL | IMN | TRUE | 36,342 | 2 | |
| 6 | 485,865 | 11 | 128,168,290 | TRA | IMN | TRUE | N/A | | |
| 6 | 507,001 | 6 | 525,794 | DUP | IMN | TRUE | 18,793 | 2 | |
| 6 | 44,227,710 | 7 | 98,922,289 | TRA | I | TRUE | N/A | | Low AF |
| 7 | 32,260,329 | 7 | 32,270,617 | DEL | IMN | TRUE | 10,288 | | |
| 7 | 69,760,460 | 7 | 69,919,119 | DEL | IMN | TRUE | 158,659 | 3 | |
| 7 | 153,902,183 | 7 | 153,912,278 | DEL | IMN | TRUE | 10,095 | | |
| 7 | 157,250,902 | 10 | 92,978,365 | TRA | IMN | TRUE | N/A | | |
| 8 | 111,029,315 | 8 | 111,042,045 | DUP | IMN | TRUE | 12,730 | 2 | |
| 9 | 10,043,022 | 9 | 36,761,238 | DEL | IMN | TRUE | 26,718,216 | | |
| 9 | 21,881,977 | 9 | 22,233,170 | DEL | IMN | TRUE | 351,193 | 2 | |
| 9 | 130,997,818 | 9 | 131,094,185 | DEL | IMN | TRUE | 96,367 | | |
| 10 | 47,550,084 | 10 | 51,470,173 | DEL | M | TRUE | 3,920,089 | 2 | |
| 11 | 35,212,288 | 11 | 35,222,587 | DEL | IMN | TRUE | 10,299 | 2 | |
| 11 | 90,022,131 | 11 | 90,059,997 | DEL | IMN | TRUE | 37,866 | | |
| 14 | 106,329,902 | 14 | 107,095,124 | DEL | M | TRUE | 765,222 | 2 | Low Illumina coverage |
| 15 | 28,122,233 | 15 | 33,764,220 | INV | IMN | TRUE | 5,641,987 | | |
| 15 | 34,228,443 | 22 | 224,600,333 | TRA | IMN | TRUE | N/A | 1 | |
| 15 | 56,408,061 | 15 | 56,470,185 | DEL | IMN | TRUE | 62,124 | | |
| 16 | 1,923,736 | 16 | 88,041,400 | DUP | IMN | TRUE | 86,117,664 | | |
| 16 | 3,213,698 | 16 | 3,892,969 | INV | IMN | TRUE | 679,271 | | |
| 16 | 3,879,149 | 16 | 3,893,262 | INV | IMN | TRUE | 14,113 | | |
| 16 | 6,417,830 | 16 | 70,481,113 | DUP | IMN | TRUE | 64,063,283 | | |
| 16 | 10,014,726 | 16 | 11,340,535 | DUP | IMN | TRUE | 1,325,809 | | |
| 16 | 17,148,805 | 16 | 77,348,079 | DEL | I | TRUE | 60,199,274 | | Low Nanopore coverage at 3' end |
| 16 | 34,300,552 | 16 | 35,187,293 | DUP | MN | TRUE | 886,741 | 2 | |
| 16 | 47,905,844 | 16 | 64,948,133 | INV | IMN | TRUE | 17,042,289 | | |
| 16 | 47,907,576 | 16 | 48,516,176 | INV | I | TRUE | 608,600 | | Low AF |
| 16 | 72,998,352 | 16 | 87,021,480 | INV | IMN | TRUE | 14,023,128 | | |
| 16 | 74,862,382 | 16 | 75,122,465 | DEL | IMN | TRUE | 260,083 | | |
| 16 | 77,300,411 | 16 | 86,927,661 | DEL | IMN | TRUE | 9,627,250 | | |
| 16 | 86,927,619 | 16 | 87,021,978 | INV | IM | TRUE | 94,359 | | |
| 17 | 3,726,234 | 17 | 5,311,468 | DEL | IMN | TRUE | 1,585,234 | | |
| 17 | 7,411,108 | 17 | 9,027,626 | DEL | IMN | TRUE | 1,616,518 | 2 | |
| 17 | 73,204,079 | 17 | 74,561,943 | DEL | IM | TRUE | 1,357,864 | | |
| 18 | 6,261,886 | 18 | 7,433,790 | DEL | IMN | TRUE | 1,171,904 | 3 | |
| 18 | 9,605,838 | 18 | 10,056,069 | DEL | IMN | TRUE | 450,231 | | |
| 19 | 5,588,926 | 19 | 5,687,770 | DEL | IMN | TRUE | 98,844 | | |
| 19 | 8,029,729 | 19 | 8,076,835 | DEL | IM | TRUE | 47,106 | 2 | |
| 19 | 10,740,031 | 19 | 10,900,300 | DEL | IMN | TRUE | 160,269 | | |
| 19 | 38,954,083 | 19 | 39,171,876 | DEL | IM | TRUE | 217,793 | | |
| 22 | 22,396,921 | 22 | 23,174,300 | INV | IMN | TRUE | 777,379 | 1 | |
| 22 | 23,150,000 | 22 | 23,174,429 | INV | IMN | TRUE | 24,429 | | |
| X | 13,641,629 | X | 13,890,199 | DEL | IMN | TRUE | 248,570 | 1 | |
| X | 154,569,879 | X | 154,951,087 | DEL | MN | TRUE | 381,208 | | |
| 1 | 19,501,813 | 4 | 114,699,596 | TRA | M | FALSE | N/A | | Present in germline |
| 1 | 31,399,988 | X | 65,505,803 | TRA | I | FALSE | N/A | | |
| 1 | 33,516,334 | 7 | 24,448,711 | TRA | M | FALSE | N/A | | |
| 1 | 75,849,140 | 12 | 55,727,218 | TRA | M | FALSE | N/A | | |
| 1 | 144,680,038 | 1 | 144,954,615 | DUP | N | FALSE | 274,577 | | Present in germline |
| 1 | 144,910,918 | 1 | 146,520,738 | DEL | I | FALSE | 1,609,820 | | |
| 1 | 145,958,228 | 9 | 140,785,678 | TRA | M | FALSE | N/A | | |
| 1 | 155,596,456 | 12 | 55,727,214 | TRA | M | FALSE | N/A | | Present in germline |
| 1 | 161,540,830 | 1 | 161,622,768 | DUP | I | FALSE | 81,938 | | |
| 1 | 168,186,186 | 1 | 182,274,315 | INV | MN | FALSE | 14,088,129 | | Present in germline |
| 1 | 174,317,572 | 7 | 138,296,609 | TRA | M | FALSE | N/A | | |
| 1 | 174,319,898 | 7 | 138,296,635 | TRA | M | FALSE | N/A | | |
| 1 | 222,660,296 | 10 | 65,571,043 | TRA | M | FALSE | N/A | | Present in germline |
| 1 | 247,286,841 | 1 | 247,337,064 | INV | MN | FALSE | 50,223 | | End of chromosome |
| 1 | 247,337,977 | 13 | 42,020,906 | TRA | M | FALSE | N/A | | |
| 2 | 38,613,641 | 7 | 108,898,305 | TRA | I | FALSE | N/A | | |
| 2 | 46,448,179 | 6 | 109,858,682 | TRA | I | FALSE | N/A | | |
| 2 | 89,160,770 | 2 | 90,122,131 | INV | I | FALSE | 961,361 | | Deletion not inversion |
| 2 | 209,033,722 | 17 | 19,995,620 | TRA | M | FALSE | N/A | | Present in germline |
| 2 | 228,820,839 | 8 | 84,795,334 | TRA | I | FALSE | N/A | | |
| 3 | 75,983,387 | 19 | 33,444,266 | TRA | M | FALSE | N/A | | Present in germline |
| 3 | 85,576,567 | 7 | 141,626,511 | TRA | M | FALSE | N/A | | Present in germline |

| | | | | | | | | |
|---|---|---|---|---|---|---|---|---|
| 3 | 111,596,344 | 3 | 111,613,565 | DEL | I | FALSE | 17,221 | |
| 3 | 118,397,667 | 12 | 187,752,105 | TRA | I | FALSE | N/A | |
| 3 | 122,426,204 | 9 | 3,707,515 | TRA | I | FALSE | N/A | |
| 3 | 122,426,228 | 9 | 3,707,648 | TRA | I | FALSE | N/A | |
| 3 | 136,217,132 | 11 | 129,112,835 | TRA | I | FALSE | N/A | |
| 3 | 137,845,369 | 19 | 53,689,190 | TRA | M | FALSE | N/A | |
| 3 | 137,845,386 | 19 | 53,691,886 | TRA | M | FALSE | N/A | |
| 3 | 137,845,386 | 7 | 1,187,649 | TRA | M | FALSE | N/A | |
| 3 | 195,348,011 | 3 | 195,476,035 | DUP | N | FALSE | 128,024 | Present in germline |
| 4 | 9,718,690 | 21 | 233,806,171 | TRA | I | FALSE | N/A | |
| 4 | 144,940,998 | 4 | 145,062,404 | DUP | I | FALSE | 121,406 | |
| 4 | 180,912,835 | 5 | 45,310,939 | TRA | I | FALSE | N/A | |
| 5 | 17,520,843 | 5 | 17,598,526 | INV | M | FALSE | 77,683 | Present in germline |
| 5 | 21,497,378 | 5 | 34,197,448 | INV | M | FALSE | 12,700,070 | Present in germline |
| 5 | 21,506,327 | 6 | 58,137,659 | TRA | M | FALSE | N/A | Start of centromere - Ns in reference |
| 5 | 39,787,755 | 9 | 94,416,754 | TRA | M | FALSE | N/A | |
| 5 | 136,854,543 | 10 | 133,083,802 | TRA | I | FALSE | N/A | |
| 5 | 141,454,222 | 13 | 82,367,712 | TRA | M | FALSE | N/A | |
| 5 | 141,456,961 | 13 | 82,367,697 | TRA | M | FALSE | N/A | |
| 5 | 156,084,716 | 10 | 27,182,398 | TRA | M | FALSE | N/A | |
| 5 | 179,060,981 | 5 | 179,082,164 | INV | M | FALSE | 21,183 | End of chromosome |
| 6 | 32,491,477 | 6 | 32,527,495 | DUP | N | FALSE | 36,018 | Present in germline - segmental duplication |
| 7 | 1,185,094 | 11 | 5,126,945 | TRA | M | FALSE | N/A | |
| 7 | 1,187,652 | 11 | 5,126,964 | TRA | M | FALSE | N/A | |
| 7 | 21,044,287 | 7 | 21,417,087 | DEL | I | FALSE | 372,800 | |
| 7 | 21,044,344 | 7 | 21,417,432 | DUP | I | FALSE | 373,088 | |
| 7 | 25,087,510 | 13 | 61,462,338 | TRA | M | FALSE | N/A | |
| 7 | 35,914,922 | 11 | 150,250,251 | TRA | I | FALSE | N/A | |
| 7 | 55,830,658 | 7 | 65,288,203 | INV | I | FALSE | 9,457,545 | Present in germline |
| 7 | 62,748,724 | 7 | 62,918,259 | INV | N | FALSE | 169,535 | Present in germline. Near centromere |
| 7 | 66,625,803 | 7 | 72,116,397 | INV | I | FALSE | 5,490,594 | |
| 7 | 71,918,049 | 7 | 72,129,584 | DEL | I | FALSE | 211,535 | |
| 7 | 81,789,164 | 10 | 60,902,931 | TRA | M | FALSE | N/A | Present in germline |
| 7 | 102,235,710 | 7 | 102,328,197 | DEL | M | FALSE | 92,487 | Present in germline |
| 7 | 142,110,010 | 7 | 142,129,906 | DEL | I | FALSE | 19,896 | |
| 7 | 143,218,861 | 7 | 143,504,793 | INV | N | FALSE | 285,932 | |
| 8 | 16,733,860 | 21 | 235,841,288 | TRA | I | FALSE | N/A | |
| 8 | 16,734,191 | 21 | 235,841,318 | TRA | I | FALSE | N/A | |
| 8 | 96,265,729 | 19 | 18,835,610 | TRA | NM | FALSE | N/A | Present in germline |
| 9 | 181,827 | 9 | 70,492,285 | DUP | M | FALSE | 70,310,458 | Present in germline |
| 9 | 200,778 | 9 | 70,835,468 | INV | N | FALSE | 70,634,690 | |
| 9 | 90,372,455 | 20 | 4,082,830 | TRA | M | FALSE | N/A | |
| 9 | 126,741,382 | 9 | 126,756,051 | INV | M | FALSE | 14,669 | Present in germline |
| 9 | 136,104,027 | 9 | 136,184,725 | INV | I | FALSE | 80,698 | |
| 10 | 38,727,633 | 18 | 114,605,803 | TRA | I | FALSE | N/A | |
| 10 | 82,293,090 | 16 | 75,503,334 | TRA | M | FALSE | N/A | |
| 10 | 82,295,349 | 16 | 75,503,316 | TRA | M | FALSE | N/A | |
| 11 | 1,915,272 | 11 | 1,961,085 | INV | N | FALSE | 45,813 | Present in germline |
| 11 | 11,268,015 | 15 | 86,440,793 | TRA | M | FALSE | N/A | |
| 11 | 11,268,778 | 15 | 86,439,512 | TRA | M | FALSE | N/A | |
| 11 | 55,365,429 | 11 | 55,445,868 | INV | M | FALSE | 80,439 | Present in germline |
| 11 | 56,853,645 | 12 | 125,801,164 | TRA | N | FALSE | N/A | Present in germline |
| 11 | 66,162,089 | 12 | 96,233,592 | TRA | M | FALSE | N/A | |
| 11 | 89,507,166 | 11 | 89,810,343 | INV | M | FALSE | 303,177 | Present in germline |
| 11 | 89,626,400 | 11 | 89,690,477 | INV | N | FALSE | 64,077 | |
| 11 | 124,110,186 | 11 | 124,121,287 | DUP | N | FALSE | 11,101 | Present in germline |
| 12 | 11,192,448 | 12 | 11,218,061 | DEL | M | FALSE | 25,613 | Present in germline |
| 12 | 59,029,090 | 12 | 70,594,878 | DEL | M | FALSE | 11,565,788 | Present in germline |
| 14 | 19,865,208 | 14 | 20,424,756 | INV | M | FALSE | 559,548 | Present in germline |
| 16 | 14,988,609 | 16 | 15,031,366 | INV | M | FALSE | 42,757 | Present in germline |
| 16 | 70,151,142 | 16 | 74,397,976 | INV | N | FALSE | 4,246,834 | Present in germline |
| 16 | 75,238,566 | 16 | 75,256,652 | INV | M | FALSE | 18,086 | Present in germline - segmental duplication |
| 19 | 8,018,972 | 19 | 8,029,672 | DUP | I | FALSE | 10,700 | Present in germline. Alu element coupled with deletion |
| 19 | 22,132,988 | 19 | 23,130,957 | DUP | I | FALSE | 997,969 | |
| 19 | 55,333,906 | 19 | 55,367,975 | DEL | I | FALSE | 34,069 | |
| 20 | 25,900,389 | 21 | 9,411,193 | TRA | M | FALSE | N/A | |
| 21 | 10,945,338 | 21 | 11,177,588 | DEL | N | FALSE | 232,250 | Present in germline |
